# DRomics, a workflow to exploit dose-response omics data in ecotoxicology

**DOI:** 10.1101/2023.02.09.527852

**Authors:** Marie Laure Delignette-Muller, Aurélie Siberchicot, Floriane Larras, Elise Billoir

## Abstract

Omics technologies has opened new possibilities to assess environmental risks and to understand the mode(s) of action of pollutants. Coupled to dose-response experimental designs, they allow a non-targeted assessment of organism responses at the molecular level along an exposure gradient. However, describing the dose-response relationships on such high-throughput data is no easy task. In a first part, we review the software available for this purpose, and their main features. We set out arguments on some statistical and modeling choices we have made while developing the R package DRomics and its positioning compared to others tools. The DRomics main analysis workflow is made available through a web interface, namely a shiny app named DRomics-shiny. Next, we present the new functionalities recently implemented. DRomics has been augmented especially to be able to handle varied omics data considering the nature of the measured signal (e.g. counts of reads in RNAseq) and the way data were collected (e.g. batch effect, situation with no experimental replicates). Another important upgrade is the development of tools to ease the biological interpretation of results. Various functions are proposed to visualize, summarize and compare the responses, for different biological groups (defined from biological annotation), optionally at different experimental levels (e.g. measurements at several omics level or in different experimental conditions). A new shiny app named DRomicsInterpreter-shiny is dedicated to the biological interpretation of results. The institutional web page https://lbbe.univ-lyon1.fr/fr/dromics gathers links to all resources related to DRomics, including the two shiny applications.

## Introduction

Dose-response (DR) modeling belongs to the toolkit of ecotoxicologists. The latter are used to this approach when working on apical endpoints (e.g. reproduction, photosynthesis). The derived sensitivity thresholds, e.g. Effective Concentrations (ECx), BenchMark Dose (BMD), are at the basis of regulatory risk assessment.

The recent years have seen the emergence of works using DR omics data (e.g. transcriptomic, proteomic, metabolomic) in ecotoxicology (Zhang *et al*., 2018). Typical DR designs, with many doses (>6), ensure a good description of the DR relationship and a robust and precise estimation of a sensitivity threshold (such as the benchmark dose – BMD) that is useful to fix regulatory thresholds (Ewald *et al*., 2022). However, such designs are less commonly used in omics studies, as tools classically used to analyse omics data are dedicated to differential expression analysis between few conditions (limma, Ritchie *et al*., 2025, DESeq2, Love *et al*., 2014, EdgeR, Robinson *et al*., 2010). The analysis of such data typically starts with a pairwise differential analysis to the control followed by an enrichment procedure to identify GO (Gene Ontology) terms or KEGG (Kyoto Encyclopedia of Genes and Genomes) biological pathways of differentially expressed items (e.g. contigs, proteins) (Dubois *et al*., 2019, Murat El Houdigui *et al*., 2019, Meier *et al*., 2020, Zhan *et al*., 2021). In a second time, some perform a DR modeling on differentially expressed items to estimate a BMD per item and summarize the sensitivity of each pathway for example by the median of BMDs of corresponding pathways (Meier *et al*., 2020, Zhan *et al*., 2021).

Studies implementing DR (multi-)omics approaches sometimes aim at a mechanistic understanding of adverse effects (Adverse Outcome Pathway perspective - AOP). They could identify potential Modes of Action of pollutants (MoAs) at the molecular level, that generally need to be validated in a second step using targeted experiments (Andersen et al. 2018). Some authors already used our R package DRomics (“Dose Response for Omics”) on transcriptomics and/or metabolomics data on different organisms and pollutants, to help the understanding of the adverse effects, by identifying most sensitive pathways (Larras *et al*., 2020; Gust *et al*., 2021; Vokuev *et al*., 2021). DRomics was also used on community transcriptomics and metabolomics data to provide insights into mechanisms of pollution-induced community tolerance (Lips *et al*., 2022; Larras *et al*., 2022). And recently, Song et al. (2023) showed, using DRomics, how DR modelling and estimation of points of departure at several omics and apical levels can be mapped to an AOP network. Those applications of DRomics especially motivated us to develop new R functions and a new shiny application to help the biological interpretation of DR modeling of omics data.

The purpose of this paper is to present the new version of the DRomics R package and its two companion interactive web applications, DRomics-shiny and DRomicsInterpreter-shiny. We first review the software available to tackle a DR analysis of high-throughput omic data and explain how DRomics distinguishes from other tools. Then, we present the various functionalities we added to DRomics from its first version published in 2018 and especially explain the way it can be used to make sense of DR (multi-)omics data in environmental risk assessment.

### DRomics original features compared to other tools dedicated to dose-response omics data

Based on literature review in toxicology and ecotoxicology, we identified five tools available for DR analysis of high-throughput omics data (Table 1). Their workflows consist of successive steps: first, the selection of items (e.g. contigs) significantly regulated along the gradient of exposure. Then, for the selected items, DR relationships are modeled and BMD are derived from these DR models. The BMD-zSD, the most often used and recommended version of the BMD, is defined as the dose that leads to a response (BMR – benchmark response) pointing a difference from the response in controls of more than z times (e.g. with z=1, EFSA, 2017) the residual standard deviation of the DR model.

**Table 1.**
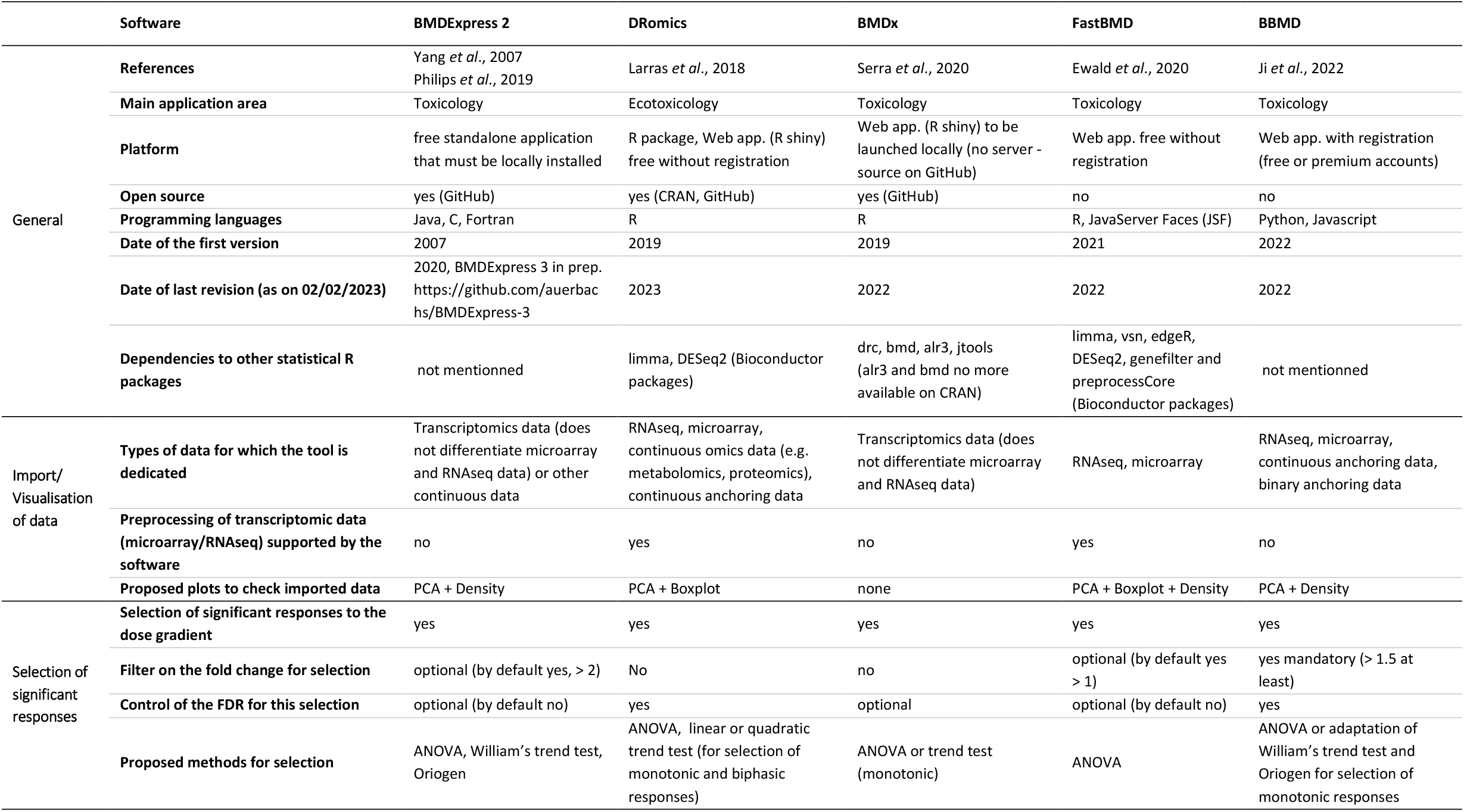
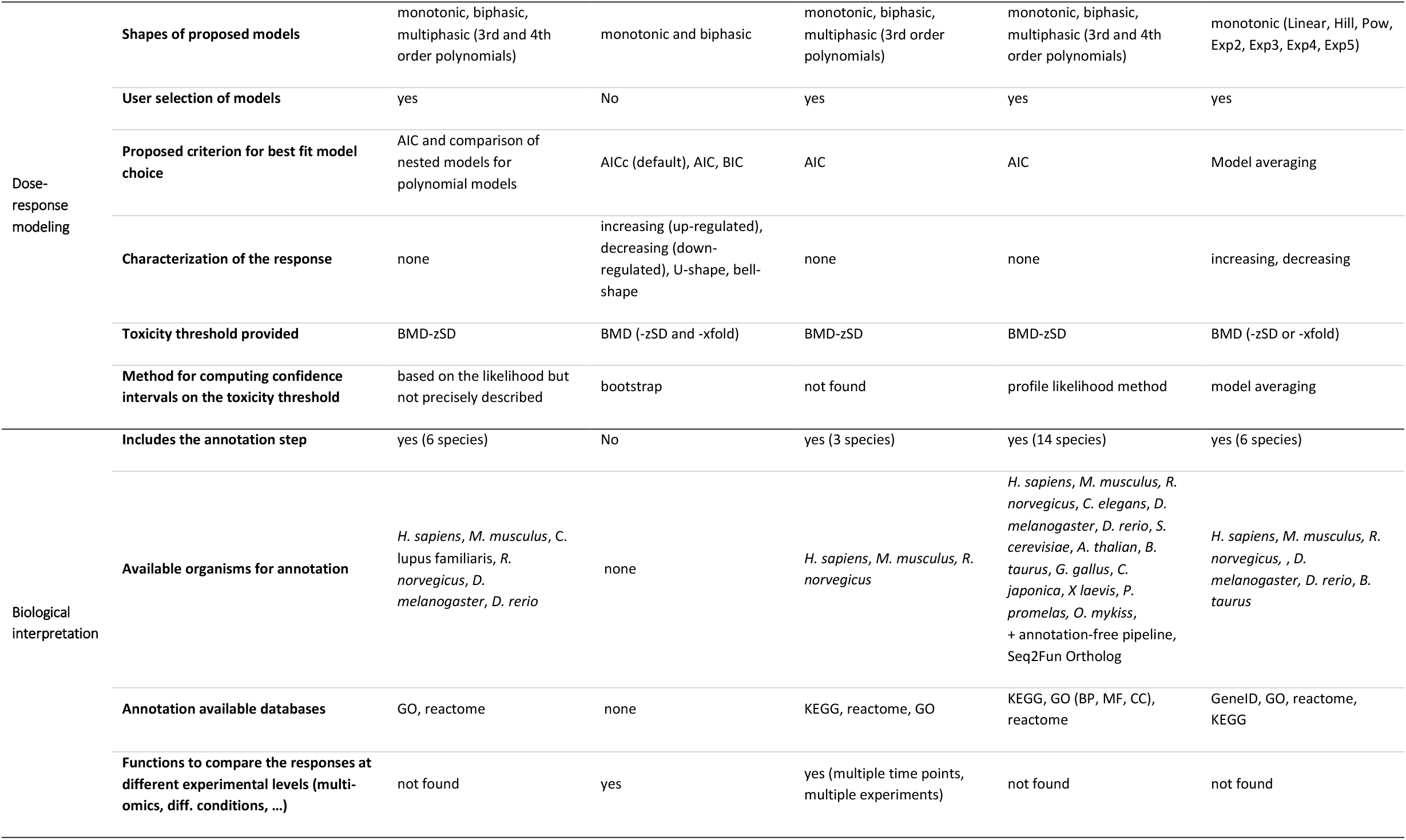
Main characteristics of the tools available for DR analysis of high-throughput omics data.

The first piece of software developed to analyze high dimensional DR data, in particular gene expression data, was BMDExpress, released in its first version in 2007 (Yang *et al*., 2007). It was since updated and augmented in BMDExpress-2 (Philips *et al*., 2019). FastBMD (Ewald *et al*., 2021) implements the same methods, as framed by the US National Toxicology Program (NTP, 2018), with the sake of being faster and friendlier to end-users than BMDExpress, and through a web-based interface. Again in the context of toxicogenomics, BBMD (Ji *et al*., 2022) has the specificity to use model averaging to account for BMD uncertainty related to the underlying DR model, and BMDx (Serra *et al*., 2020) allows a comparison of BMD values of a transcriptomics experiment at different time points or from different experiments. We developed DRomics in 2018 (Larras *et al*., 2018) for DR analysis of any omics data (e.g. transcriptomics, metabolomics), including non-sequenced organisms or communities (meta-omics), biological models commonly used in the field of ecotoxicology. The main characteristics and functionalities of those five tools are summarized in Table 1.

DRomics is the only tool which is composed of both an R package and a free web application (see Table 1). We chose to develop it in the R language among others i) to facilitate the interoperability with the Bioconductor packages that implement state-of-the-art bioinformatics methods and ii) to add a shiny application (interactive web app. straight from R) that can be launched both from the package and from a server freely accessible without registration. The companion DRomics-shiny application was thought for users who do not want to work in the R environment, but also to help new users to take the package in hand. For that purpose, the R code of the whole performed analysis is provided in the last page of the shiny application. While developing DRomics, we were cautious to limit as much as possible the dependencies to other statistical tools and to only depend on well-maintained R packages. Notice that the installation of BMDx is currently impossible because of dependencies to non-maintained R packages.

DRomics was at the root designed to be able to analyze data from typical DR design, favoring the number of doses over the number of replicates per dose, or even for datasets with no experimental replicates. This situation of DR approach with no replicates is met in some field studies (one dose per sample) and in some screening studies as illustrated in Rollin et al. (2023). This is the reason why, for the selection of significantly responsive items, we did not use classical methods based on comparison of means at different doses, which cannot be applied without replicates, such as one-way ANOVA, William’s trend test (Williams, 1971) or the ORIOGEN method (Peddada *et al*., 2003). Instead we implemented methods based on the fit of a linear or quadratic model to the data (as coarse approximation of the observed trend) using the ranks of observed doses as an independent variable (for a better robustness to the repartition of the tested/observed doses that is sometimes more regular on a log scale). Those two original methods proposed in DRomics (in addition to the classical ANOVA method) were inspired by the trend tests proposed by Tukey et al. (1985) and do not require replicates. They were implemented using robust empirical Bayesian functions provided by DESeq2 (for RNAseq data, Love *et al*., 2014) or by limma (for other omic data, Ritchie *et al*., 2015). To control the false discovery rate (FDR), DRomics implements a mandatory use of the Benjamini-Hochberg correction/adjustment. Last, we decided not to apply a fold-change filter, in order to keep weak signals if they are significant, and because fold change is difficult to define for data with no replicate. DRomics proposes as a default selection method the quadratic trend test which can detect both monotonic and biphasic responses, and is far more efficient than the classical ANOVA-type test (Larras *et al*., 2018) when the number of replicates is low.

For the DR modeling of selected items, the tools developed in the field of toxicology (Table 1) use only monotonic models (case of BBMD) or the NTP models (NTP 2018, case of BMDExpress, FastBMD and BMDx). Among the NTP models, only polynomial models can describe non-monotonic responses that are commonly occurring in omics dose-response data (Smetanova *et al*., 2015, Larras *et al*., 2018). Polynomial models of degree 3 and 4 that are proposed by all the tools using the NTP models (Table 1) are no longer recommended by the NTP as long as they cannot be constrained to change direction only once (NTP 2018). So biphasic responses can only be described by a parabole using NTP models, which does not offer a great flexibility. In the field of ecotoxicology, more flexible biphasic models were used to model biphasic DR data based on the Gaussian model (Gundel *et al*., 2012; Smetanova *et al*., 2015). As one of our main objectives, while developing DRomics, was to be able to select, model and characterize all types of monotonic and biphasic responses, we defined our own model family, especially including original flexible biphasic models, the Gauss-probit and logGauss-probit models (for a complete description see the package vignette https://cran.r-project.org/web/packages/DRomics/vignettes/DRomics_vignette.html#models or Larras *et al*., 2018). BMDExpress and DRomics results were compared in the case study of a microarray dataset, as reported in the supporting information of Larras et al. (2018). DRomics models were shown to give a better description of data (smaller AIC values) and more repeatable and conservative BMD estimations for biphasic responses. Unlike the other tools (Table 1), DRomics does not only provide a BMD estimation, but also a characterization of the DR response in four classes (increasing, decreasing, U-shape, bell-shape) which we thought may be of great interest in an AOP perspective, when the DR analysis not only aims at defining BMD values but also at making sense of DR (multi-)omics data in environmental risk assessment.

### New features in the DRomics DR modeling workflow

Figure 1 maps the DRomics workflow and the functionalities offered to explore the results. First developed for microarray data, DRomics is now able to handle RNAseq data, metabolomic or other continuous omic data (e.g. proteomics data) or even continuous non omic data (e.g. growth data) that could be used for phenotypical anchoring (respectively imported using RNAseqdata(), continuousomicdata() and continuousanchoringdata() functions). The same modeling workflow, choosing the best fit models among our complete family of models (as described previously), was declined for each type of data (e.g. intensities, counts) to ensure the comparability of results (e.g. transcriptomics vs. metabolomics, transcriptomics vs. anchoring). Writing in the R language ensures the interoperability with functions of the Bioconductor packages. Thus, functions of the DESeq2 and limma packages are internally called within the package to normalize and/or transform omic data and to implement the trend tests for selecting significantly responsive items. The call to additional R functions can be added for example to correct omics data for a potential batch effect (see an example using ComBat-seq in the package vignette: https://cran.r-project.org/web/packages/DRomics/vignettes/DRomics_vignette.html#batcheffect).

**Figure 1.**
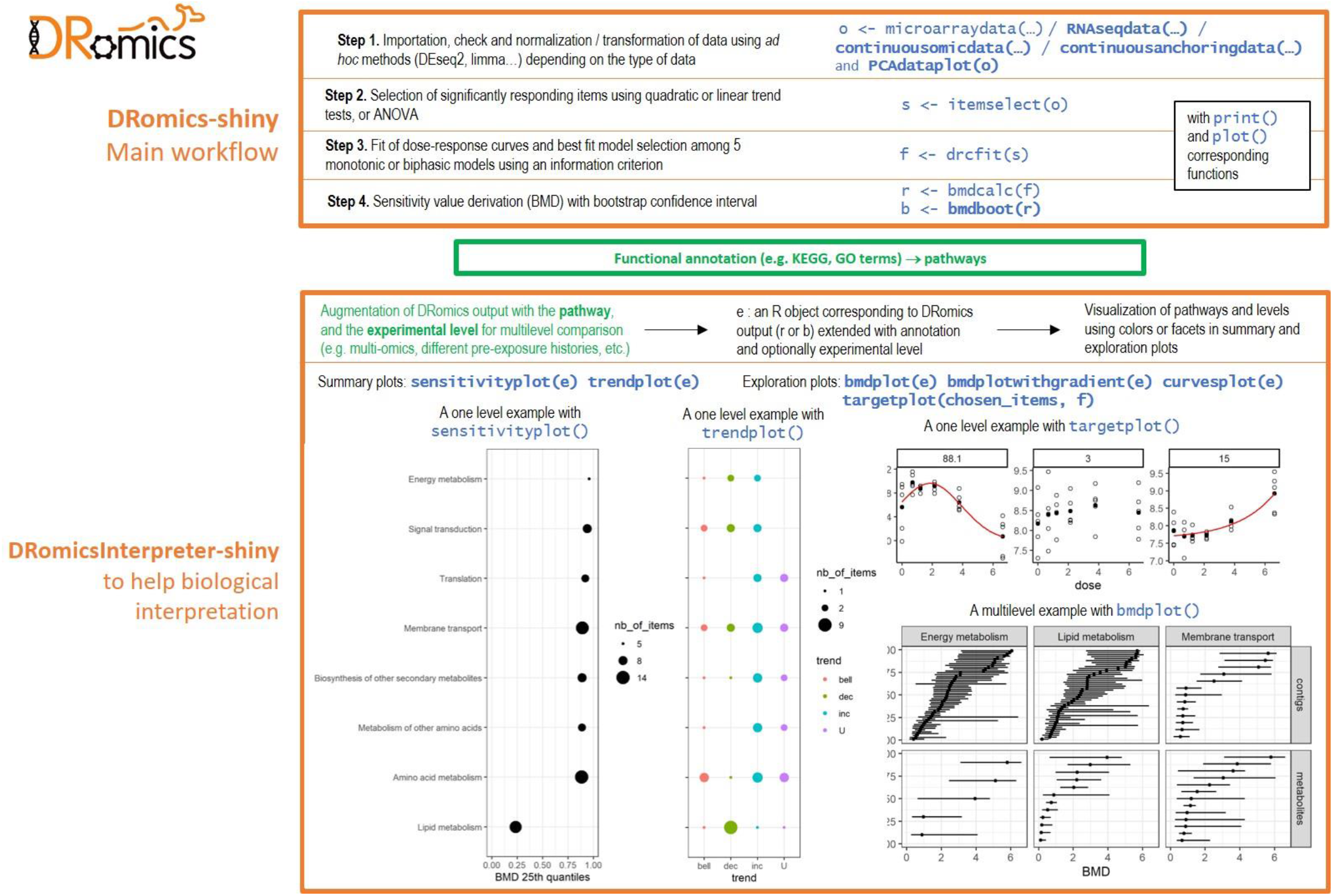
Diagram of the DRomics functionalities and of the perimeter of associated tools. Bold font indicates new functions added to the DRomics R package since its first version.

Various functions were also added to the package (highlighted in bold type in Figure 1) to respond to user requests. Among them we can cite PCAdataplot() for a visualization of data after the importation step and detection of potential outlier samples or batch effect, targetplot() to visualize the response of targeted items whatever they are selected or not in the DRomics workflow, bmdboot() and bmdplot() to compute and visualize confidence intervals on BMD values using bootstrap.

Moreover, we performed modifications in the modeling workflow to ensure a better robustness of results on data with a low number of doses. For example, we changed the default information criterion used for best model selection from the AIC to the AICc, as recommended by Burnham and Anderson (2004), and limited the set of models for weak designs with few doses (4 or 5). Despite this care one should favor optimal dose-response designs with more doses (at least 6-7, and never less than 4) and less (or no) replicates as recommended by statisticians in toxicology (Moore and Caux, 1997; Ritz, 2010; Larras *et al*., 2018; Ewald *et al*., 2022).

### New functions and the new shiny application to help biological interpretation

In toxicology, while working on sequenced and well-annotated organisms, items (e.g. genes) can be functionally annotated prior to DR analysis. BMDExpress integrates this annotation step for 6 model species on the basis of GO or reactome databases, and FastBMD for 14 model species on the basis of GeneID, GO, Reactome or KEGG databases (see Table 1). Such an annotation of all the items whatever they respond or not to the dose gradient exposition, enables the classical enrichment analysis (Wu *et al*., 2021). This analysis consists in highlighting gene sets/pathways which are the most overrepresented among the responding ones.

In ecotoxicology, one often works on non-model organisms or even on samples from environmental communities (freshwater biofilm for example – Creusot *et al*., 2021; Larras et al. 2022, Lips *et al*., 2022).

This implies the need for the user to retrieve and manage its own annotation, which is a challenging task, especially for RNAseq experiments for which the number of measured contigs may be huge (several millions of contigs). We thus considered that this annotation step could be done after the selection/modeling workflow, to reduce the number of items to annotate, and so the difficulty of this task. Due to the great diversity of annotation pipelines that can be developed for such non-model organisms, we did not include an annotation step in DRomics. However, and to support the interpretation of the workflow results in view of a biological annotation provided by the user, we recently developed new functions and a second shiny application (named DRomicsInterpreter-shiny) (Figure 1).

After augmenting the DRomics output with information about the functional role of items, the graphical representations offered by DRomics give a new insight into the results. Further, provided a common annotation system is used at different biological scales under study, DRomics can be used to compare the response at various levels. Figures 2 and 3 give some illustrations on an example with two molecular levels from Larras et al. (2020): transcriptomics and metabolomics responses of *Scenedesmus vacuolatus* to triclosan exposure. Various functions are proposed to visualize/summarize/compare the responses, for the different biological groups (defined from biological annotation), optionally at the different experimental levels.

**Figure 2.**
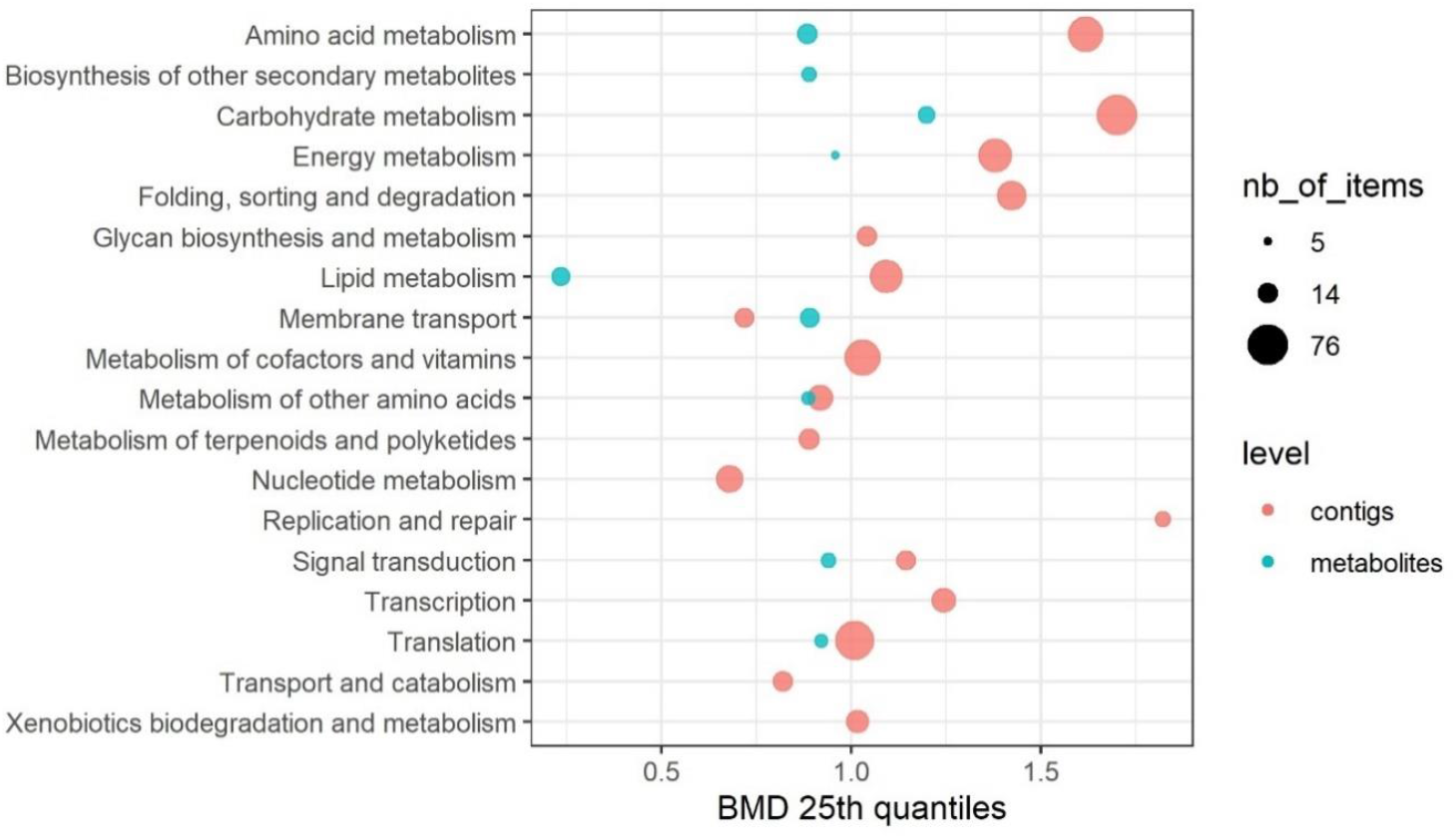
Illustration of the use of the function sensitivityplot() here to summarize the sensitivity of the responding KEGG pathways at two molecular levels using transcriptomic (contigs) and metabolomic (metabolites) data published in Larras et al. (2020) on *Scenedesmus vacuolatus* exposed to triclosan.

**Figure 3.**
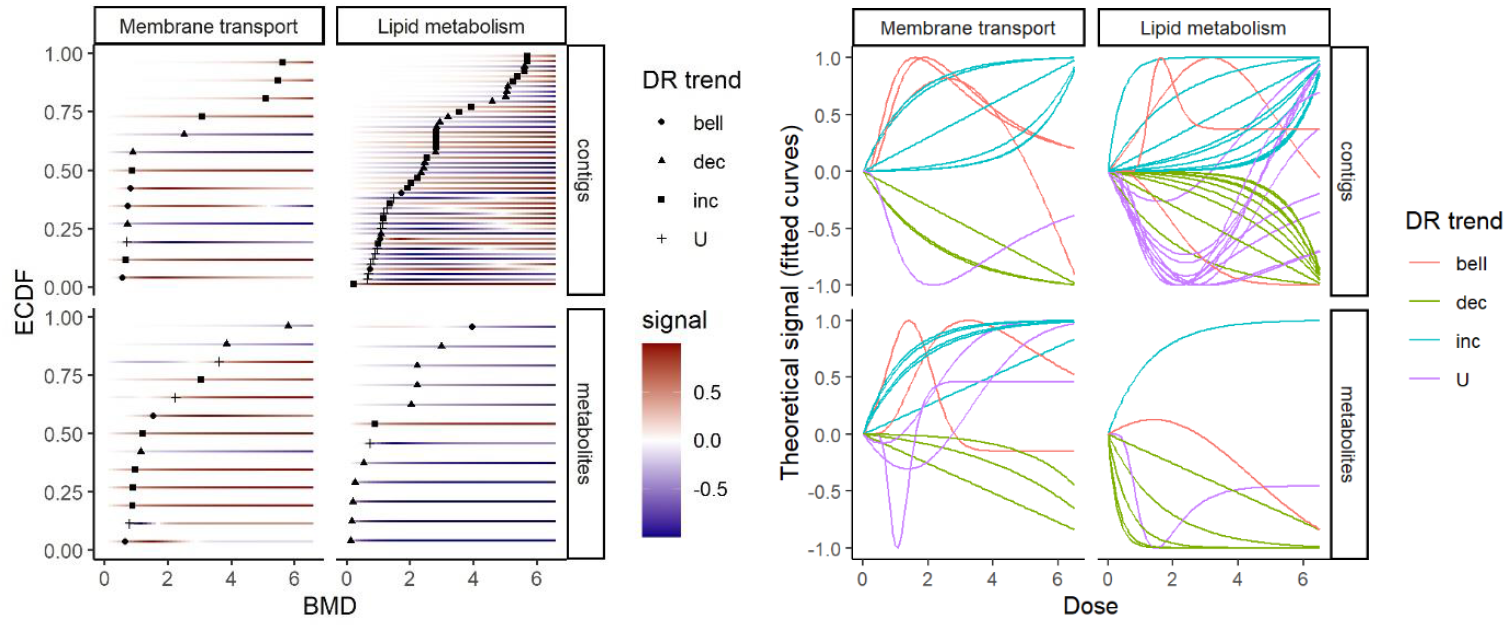
Summary of the dose response (DR) curves of all contigs and metabolites corresponding to two specific KEGG pathways: «membrane transport» and «lipid metabolism» (from data published in Larras et al. (2020) on Scenedesmus vacuolatus exposed to triclosan) on the left using the bmdplotwithgradient() function and on the right using the curvesplot() fuction.

- the trends (increasing, decreasing, U-shape or bell-shape) of the DR curves (using the trendplot() function, see a one-level example on Figure 1),
- the group/pathway-level sensitivity calculated as a quantile of BMD values of the group (using the sensitivityplot() function, see an one-level example on Figure 1 and a multi-level example on Figure 2),
- for selected groups/pathways, all the BMD values with their confidence intervals (using the bmdplot() function, see a multilevel-level example on Figure 1, right part)
- for selected groups/pathways, all the BMD values with the corresponding DR curve signal coded as a color gradient (using the bmdplotwithgradient() function, see a multi-level example on Figure 3, left part).
- for selected groups/pathways, all the DR curves represented as curves (using the curvesplot() function, see a multi-level example on Figure 3, right part),

Those functions can also be used to compare the response at one molecular level but measured under different experimental conditions (different time points, different pre-exposure scenarios, in vitro/in vivo, etc.). The selectgroups() function was also developed to help the user to focus its interpretation on the most represented and/or the most sensitive biological groups (see an example in the package vignette: https://cran.r-project.org/web/packages/DRomics/vignettes/DRomics_vignette.html#selectgroups).

### Perspectives

In the report of the National Toxicology Program (NTP, 2018) model averaging is mentioned as an interesting feature to make the BMD estimate less dependent of the choice of the best DR model. BBMD (Ji *et al*., 2022) and the next version of BMDExpress in preparation (BMDExpress-3 – https://github.com/auerbachs/BMDExpress-3/) include a Bayesian model averaging procedure. However, we do not plan to implement model averaging in DRomics because the BMD estimation is not our only purpose. We also want to characterize each response by its trend, which is itself dependent of the model choice and not averageable. Instead, we plan to add an alternative to our current bootstrap procedure, enabling the fit of a different model at each bootstrap iteration, to be able to pass the uncertainty due to the model choice both on the BMD uncertainty and on the trend uncertainty.

Concerning the modeling workflow, so far, we added specific functionalities to analyse continuous anchoring data, essentially to prevent comparisons of BMD values at different biological scales but with BMD obtained from different analysis workflows, which could induce a bias. As anchoring data may be non-continuous, such as dichotomous survival data, or reproduction data reported as number of offspring per individual-day (Delignette-Muller *et al*., 2014), we plan new developments in DRomics to be able to analyse those data properly taking into account their nature.

Concerning our recent development of functions to help the biological interpretation of DRomics results, we plan to enlarge the range of the methods proposed, by imaging new plots and summaries to explore and characterize the responses and their diversity within a biological group/pathway.

## Conclusion

The development of DRomics has been driven by the demands of ecotoxicologists to help make full sense of their dose-response (multi-)omics studies. Hence, DRomics was augmented to provide a common workflow to handle (meta-)transcriptomics, proteomics, metabolomics and/or anchoring data. This creates the foundations for a proper comparison of responses at different omics levels (and anchoring endpoints) and a mechanistic understanding in an AOP perspective. Along with functional annotations, DRomics outputs (response trends, BMD, etc.) can now be processed using a series of graphical functions thought to help their biological interpretation at the metabolic pathway level. The comparison is made easy (i) of different measurements, for instance transcripto- and metabolomics, (ii) of different biological materials, for instance communities with/without pre-exposure history, or (iii) of experimental settings, for instance successive timepoints or different temperatures. Moreover, two special cases have been addressed: experiments with a batch effect and designs with no replicates. DRomics future direction and evolutions depend on upcoming challenges and needs brought by (eco)toxicologists.

## Acknowledgements

The authors thank colleagues and students whose requests and/or discussions contributed to the recent evolution of DRomics, especially Olivier Armant, Ellis Franklin, Sandrine Frelon, Sophie Prud’homme, Mechthild Schmitt-Jansen and Philippe Veber. We also want to thank the referees for their helpful suggestions that contributed to the improvement this manuscript.

## Data, scripts, code, and supplementary information availability

Scripts and code are available online: DRomics is written in R, freely available from CRAN and encompasses two shiny applications (https://cran.r-project.org/web/packages/DRomics/). Those shiny applications are also freely available online. The homepage https://lbbe.univ-lyon1.fr/fr/dromics gathers links to all resources related to DRomics, e.g. the two shiny applications, a cheat sheet and a complete vignette to start with DRomics. The github https://aursiber.github.io/DRomics/ hosts DRomics stable as well as development versions. Data used in this paper are embedded in the DRomics package.

## Conflict of interest disclosure

The authors declare that they comply with the PCI rule of having no financial conflicts of interest in relation to the content of the article.

## Funding

This work has been financially supported by the Graduate School H2O’Lyon (ANR-17-EURE-0018) of Université de Lyon (UdL), within the program “Investissements d’Avenir” operated by the French National Research Agency (ANR)”, by the “Ecosphère continentale et côtière” (EC2CO) interdisciplinary program from the Centre National de la Recherche Scientifique (CNRS, France), as parts of the DROMADERE and the ECORA projects, and by the “Agence Nationale de la Recherche” as a part of the Chroco project (ANR-21-CE34-0003).

## Notes

https://lbbe.univ-lyon1.fr/fr/dromics

### Competing Interest Statement

The authors have declared no competing interest.

### Summary of Updates

We just added on the first page of this new version the PCI Ecotox Env Chem badge, with an hyperlink pointing to the recommendation of this article by PCI Ecotox Env Chem.

